# Defining a hypoxic exposure paradigm for modelling preeclampsia-associated syncytiotrophoblast dysfunction in human blastocyst-derived placental stem cells

**DOI:** 10.64898/2026.11.10.765732

**Authors:** Jenny S. Feeney, Cristina Monaco, Tina Podinic, Samantha L. Wilson, Sandeep Raha

## Abstract

**Introduction:** Preeclampsia (PE) is a hypertensive pregnancy disorder in which placental dysfunction is thought to play a central role. Although hypoxia is widely used to model PE-associated syncytiotrophoblast (ST) dysfunction in vitro, considerable variability in oxygen control conditions and hypoxic exposure paradigms has limited standardization across studies. This study aimed to establish physiologically relevant oxygen control conditions and identify a hypoxic exposure paradigm that consistently recapitulates characteristic features of PE-associated ST dysfunction.

**Methods:** Human blastocyst-derived trophoblast stem cells were differentiated into STs and exposed to physiological (8% O_2_), atmospheric (21% O_2_), or hypoxic (1% O_2_) conditions for 24 or 48 h. Angiogenic and apoptotic marker expression, cell death, and mitochondrial stress were assessed.

**Results:** Compared with atmospheric oxygen, 8% O_2_ reduced cell death and more closely reflected physiological ST conditions. Acute hypoxia (24 h) induced a robust PE-like phenotype in STs characterized by induction of anti-angiogenic markers, sFlt-1 and *sEng*, reduced expression of the pro-angiogenic factor PlGF, and increased cell death, apoptotic signalling and mitochondrial stress. Notably, induction of several of these markers was attenuated or reversed following prolonged hypoxic exposure.

**Conclusions:** These findings identify 8% O_2_ as a physiologically relevant control condition for modelling PE-associated ST dysfunction relative to 21% O_2_, and demonstrate that 24 h of severe hypoxia more consistently reproduces key molecular and cellular features of PE than prolonged hypoxia. This work establishes a framework for modelling hypoxia-induced ST dysfunction while highlighting the importance of oxygen control conditions and hypoxic exposure duration when modeling PE in vitro.

## 1. Introduction

Preeclampsia (PE) is a hypertensive pregnancy disorder that affects approximately 2-8% of pregnancies worldwide [1]. It is diagnosed by new-onset hypertension occurring after 20 weeks of gestation, accompanied by at least one of the following: proteinuria, uteroplacental dysfunction, or other maternal organ dysfunction [2,3]. PE is commonly classified according to the timing of symptom presentation, with early-onset preeclampsia (EOPE) occurring prior to 34 weeks’ gestation, and late-onset preeclampsia (LOPE) occurring at or after 34 weeks’ gestation [4]. Accounting for over 500,000 fetal and 70,000 maternal deaths per year worldwide, PE remains a leading cause of maternal and fetal morbidity and mortality [2,5]. Despite its clinical burden, the mechanisms underlying PE remain largely unclear, and treatment options remain very limited, with delivery of the placenta remaining the only definitive cure [6–8]. Identifying the specific mechanisms underlying PE is therefore critical to the development of targeted therapies that may improve maternal and fetal outcomes.

Placental dysfunction is thought to play a central role in the development and progression of PE [9]. The classical two-stage model proposes that incomplete spiral artery remodelling leads to placental malperfusion and reduced oxygen availability, resulting in placental stress and syncytiotrophoblast (ST) dysfunction that ultimately contribute to the maternal clinical manifestations of the disorder [8,10]. While deficient spiral artery remodelling is more strongly associated with the EOPE subtype, LOPE is also thought to involve placental malperfusion, resulting from mechanical constraints on placental growth that ultimately lead to similar placental stress and downstream dysfunction [10]. The resulting reduction in oxygen availability (placental hypoxia) is therefore thought to contribute to ST dysfunction in PE and has been associated with oxidative stress, mitochondrial dysfunction, trophoblast cell death, and dysregulated secretion of trophoblast-derived anti-angiogenic and inflammatory factors [3,11]. Together, these changes contribute to maternal vascular dysfunction and the clinical manifestations of the disorder [11–13].

Accurately reproducing the placental microenvironment and associated ST dysfunction observed in PE experimentally remains challenging. Human placental explants are difficult to obtain due to limited tissue availability, and animal models incompletely recapitulate human placental development or PE pathology [14]. Furthermore, PE occurs primarily in humans and only rarely in a limited number of non-human primate species [15]. In vitro trophoblast models are therefore widely used to investigate the cellular mechanisms underlying PE-associated placental dysfunction [14]. However, the physiological relevance of these models depends not only on the selected trophoblast model, but also on the experimental conditions used to define both healthy and pathological states.

Hypoxia is commonly used in vitro to mimic the placental malperfusion and reduced oxygen availability associated with PE, with severe hypoxia (typically ∼1% O_2_) frequently employed to induce PE-associated trophoblast stress [14,16–18]. However, considerable variability exists among studies in the oxygen concentrations used, the timing of exposure relative to trophoblast differentiation, and the duration of hypoxia. The lack of standardization limits comparison across studies and makes it difficult to determine which exposure paradigm more consistently reproduces PE-associated ST dysfunction.

One major concern is the selection of the control oxygen condition, whereby atmospheric oxygen (21% O_2_) is frequently used as a physiological control, despite greatly exceeding the ∼8% oxygen environment of the healthy second-trimester placenta [16,19,20]. Atmospheric oxygen therefore represents a relative hyperoxic condition for placental cells, and could independently induce cellular stress, ultimately confounding the interpretation of hypoxia-induced effects [16,21]. Moreover, it remains unclear whether prolonged hypoxic exposure amplifies the pathological changes following acute hypoxia or instead induces distinct cellular adaptations [16,18,22].

Many studies rely on transformed trophoblast cell lines that incompletely reflect primary trophoblast biology [14,23]. In contrast, human blastocyst-derived trophoblast stem cells can be differentiated into specialized trophoblast lineages, including STs, and more closely recapitulate key transcriptional and functional features of human trophoblasts [24,25]. These cells may therefore offer a more physiologically relevant system for defining appropriate oxygen conditions and modelling PE-associated trophoblast dysfunction.

To address these methodological gaps, the present study first compared atmospheric oxygen with oxygen levels representative of the healthy second-trimester placenta to establish a physiological reference condition for human blastocyst-derived STs. Using this condition as the comparator, we then assessed acute and prolonged exposure to severe hypoxia to identify the paradigm that most consistently reproduces key features of PE-associated ST dysfunction, including angiogenic imbalance, cell death, apoptotic signalling, and mitochondrial stress.

## 2. Materials & Methods

### 2.1. Cell Culture

Human blastocyst-derived trophoblast stem cells (RCB-4940, Riken BRC Cell Bank, bTS5) were passaged and used at passage 33 or earlier for all experiments. Cells were cultured in medium consisting of Dulbecco’s Modified Eagle Medium/Nutrient Mixture F-12 (DMEM/F-12 GlutaMAX, Fisher Scientific, 10565042), supplemented with 1% ITS (insulin-transferrin-selenium, Fisher Scientific, 41400045), 0.5% penicillin-streptomycin (Pen/Strep, Fisher Scientific, 15140122), 0.3% bovine serum albumin (BSA, Sigma-Aldrich, A9205), 0.2% fetal bovine serum (FBS, ThermoFisher, 16141079), 1.5 µg/mL L-ascorbic acid (Sigma-Aldrich, A5960), 50 ng/mL human recombinant epidermal growth factor (EGF, Stemcell Technologies, 78006), 0.8 mM valproic acid (VPA, Stemcell Technologies, 100-1053), 0.1 mM 2-mercaptoethanol (Sigma-Aldrich, 8057401000), 5 µM Y-27632 (Stemcell Technologies, 72304), 2 µM CHIR99021 (Stemcell Technologies, 75052), 1 µM SB431542 (Stemcell Technologies, 72234), and 0.5 µM A83-01 (Stemcell Technologies, 72022) [25]. Tissue culture dishes were coated with 10 µg/mL Collagen IV (Col IV, Sigma-Aldrich, C7521) in Dulbecco’s phosphate-buffered saline (DPBS) and incubated for a minimum of 3 h at 37°C before plating. Maintenance cultured cells were incubated at 37°C, 5% CO_2_, and 95% atmospheric air conditions. Media was replaced every 48 h and cells were passaged once they reached 70-80% confluency by washing with DPBS, detaching through a 10-min incubation with TrypLE (Fisher Scientific, 12605010), and splitting at a 1:3 ratio.

### 2.2. Syncytiotrophoblast Differentiation and Hypoxic Exposure

For all experiments, cells were passaged and immediately seeded in differentiation medium at a density of 3.3 × 10^4^ cells/cm^2^ in multi-well plates pre-coated with 10 µg/mL Collagen IV in DPBS. Differentiation medium consisted of DMEM/F-12 GlutaMAX, supplemented with 4% KnockOut Serum Replacement (KSR, Fisher Scientific, 10828028), 1% ITS, 0.5% Pen/Strep, 0.3% BSA, 0.1 mM 2-mercaptoethanol, 2.5 µM Y-27632, and 2 µM forskolin (FSK, Sigma-Aldrich, 344270) [25]. Cells were differentiated and incubated at 37°C, 5% CO_2_, and 95% atmospheric air for 96 h to allow for complete syncytialization to occur, with differentiation medium replacements every 24 h [26]. Following 96 h of syncytialization, cells were either transferred to 1% O_2_ (hypoxic) or 8% O_2_ (physiological) conditions, or maintained at 21% O_2_ (atmospheric) conditions at 37 °C and 5% CO_2_ for an additional 24 h (acute exposure) or 48 h (prolonged exposure). All low-oxygen cell culture (1% O_2_ and 8% O_2_) was carried out in the BioSpherix^®^ Xvivo environmental chamber (BioSpherix) to allow for handling under constant low oxygen levels, with media pre-equilibrated for 24 h at the appropriate oxygen concentrations.

### 2.3. RNA Extraction & RT-qPCR

Following 24 or 48 h of exposure to varying oxygen conditions, RCB 4940 cells were harvested by lysing with 500µL of cold TRIzol^TM^ reagent (ThermoFisher, 15596026) and transferred to Eppendorf tubes. Tubes were vortexed for 20 s followed by a 10-min centrifugation at 12,000g. A spin-column based method using the Direct-Zol^TM^ RNA MiniPrep Kit (Zymo Research, R2050) was used for total RNA extraction according to the manufacturer’s protocol. Total RNA concentration and purity ratios were assessed using a NanoPhotometer^®^ N60 spectrophotometer (Implen) to ensure sufficient RNA yield and sample purity. 250 ng or 500 ng of RNA (depending on the experiment) was then converted into cDNA using the Applied Biosystems^TM^ High-Capacity cDNA Reverse Transcription Kit (Applied Biosystems, 4368814) according to the manufacturer’s protocol. mRNA expression was measured through quantitative real-time PCR (RT-qPCR) using GB-Amp^TM^ SYBR Green qPCR Mix (GeneBio, P2092) and performed using the CFX384 Touch^TM^ Real-Time PCR Detection System (Bio-Rad Laboratories). The delta-delta Ct (2^-ΔΔCt^) method was used in quantifying the fold-change in mRNA expression levels. Analysis of fold-change was normalized to *β-actin*, a housekeeping gene, and vehicle control cells exposed to 8% oxygen.

### 2.4. BCA Assay

The Pierce^TM^ BCA Protein Assay Kit (ThermoFisher, 23227) was used to determine total protein concentrations. 0-2000μg/mL Bovine Serum Albumin (BSA) was used as a standard and measured at an absorbance of 562 nm in the Tecan Spark^®^ Multimode Microplate Reader.

### 2.5. SDS-Page and Western Blotting

RCB 4940 cells were seeded in multi-well plates and differentiated as described above. Following 24 or 48 h of exposure to varying oxygen conditions, cells were harvested using radioimmunoprecipitation assay (RIPA) lysis buffer with protease and phosphatase inhibitors (Roche, 4693159001 and 04906837001). Samples were prepared from total protein extract, Laemelli Sample Buffer (4X or 6X) and β-mercaptoethanol. Equal amounts of protein (10 or 15 µg/sample, depending on the experiment) were loaded on 15-well 4-15% gradient or 10% Mini-PROTEAN TGX Stain-Free gels and separated by gel electrophoresis. Following electrophoresis, stain-free gels were imaged using a ChemiDoc^TM^ Imaging System (Bio-Rad Laboratories) prior to protein transfer. Protein gels were then transferred using the Trans-Blot Turbo Transfer System (Bio-Rad Laboratories) onto polyvinylidene fluoride (PVDF) membranes. Membranes were then blocked in 1X Tris-buffered saline-Tween^®^ 20 (TBS-T) with 5% BSA overnight at 4°C, and probed using anti-HIF-1α (Abcam, ab179483, rabbit monoclonal, 1:1000) and anti-p53 (Santa Cruz Biotechnology, sc-6243, rabbit polyclonal, 1:1000) for 2 h at room temperature in the blocking solution. Membranes were then washed in TBS-T and incubated with goat anti-rabbit (Cell Signalling Technology, 1:5000) secondary antibody for 1 h at room temperature for immunodetection of the primary antibodies. Following TBS-T washes, bands were visualized with Clarity Max Western enzyme-linked chemiluminescence (ECL) substrate (Bio-Rad Laboratories, 1705062) and imaged using the Bio-Rad ChemiDoc Imaging System. Band intensities were quantified using Bio-Rad Version 6.1 Image Lab software and normalized to total protein content of the stain-free membranes.

### 2.6. Enzyme-Linked Immunosorbent Assay (ELISA)

RCB 4940 cells were seeded in multi-well plates and differentiated as described above. Following 24 or 48 h of exposure to varying oxygen conditions, cell culture medium was collected and transferred to Eppendorf tubes. Tubes were then centrifuged for 10 min at 300g and 4°C and the supernatant was collected ad used to quantify soluble protein expression of Flt-1 and PlGF was then quantified using the Human VEGFR1/Flt-1 and PLGF Quantikine^TM^ ELISA Kits (R&D Systems, DVR100C & DPG00) according to the manufacturer’s protocols. Briefly, 100 µL of each sample, diluted as necessary, was added in duplicates into antibody-coated wells and incubated for 2 h at room temperature. Wells were then washed four times with Wash Buffer prior to the addition of horseradish peroxidase-conjugated antibodies for a 1-or 2-h incubation. Washes were then repeated and Substrate Solution was added for a 30-min incubation in the dark. Stop solution was added to each well and plates were measured at an absorbance of 450 nm and 570 nm (for optical imperfection correction) in the Tecan Spark^®^ Multimode Microplate Reader.

### 2.7. DCFDA (2′,7′-Dichlorofluorescin Diacetate) Assay

RCB 4940 cells were seeded in black-walled, clear-bottom 96-well microplates and differentiated as described above. Levels of reactive oxygen species (ROS) were detected using the DCFDA Cellular ROS Detection Assay Kit (Abcam, ab113851) according to the manufacturer’s protocol. Briefly, following 24 or 48 h of exposure to varying oxygen conditions, cells were washed with 1X Buffer, then incubated with 5 µM DCFDA for 45 min in the dark in the appropriate oxygen condition at 37 °C. Fluorescence was quantified at an excitation/emission of 485/535nm using the Tecan Spark^®^ Multimode Microplate Reader, and readings were normalized to total protein content using the Pierce^TM^ BCA Protein Assay Kit.

### 2.8. Annexin V-FITC/ Propidium Iodide Staining

Annexin V-FITC/ Propidium Iodide (PI) staining was used to measure apoptosis and necrosis using an Annexin V-FITC Apoptosis Detection Kit (Abcam, ab14085) following the manufacturer’s instructions. Briefly, cells were seeded on 4-well culture slides pre-coated with Poly D Lysine (0.1mg/mL) for 4 h at room temperature, before pre-coating with 10 µg/mL Collagen IV (for 3 h at 37°C) at a cell density of 50,000 cells/well, and differentiated as described above. Following 24 or 48 h of exposure to varying oxygen conditions, cells were washed in 1X Binding Buffer II and incubated with 1:100 Annexin V-FITC and 1:100 PI in 1X Binding Buffer II for 15 min at room temperature in the dark. Cells were then washed in 1X Binding Buffer II and fixed with 2% formaldehyde for 10 min. Slides were mounted with DAPI-containing medium and left to dry for 2 h in the dark. Slides were then imaged on 20X objective with 1 s exposure time on DAPI, FITC, and TRITC channels using the Nikon ECLIPSE Ti2 Inverted Microscope, and quantified using Fiji software. Annexin V-positive cells were quantified as a measure of apoptosis, PI-positive cells were quantified as a measure of necrosis (reflecting loss of plasma membrane integrity), and overall cell death was quantified as the combined Annexin V and PI fluorescence signal normalized to the total number of DAPI-positive nuclei.

### 2.9. Statistical Analyses

Statistical analyses were performed using GraphPad Prism Version 11 software. All experiments were performed using four to six biological replicates. Outliers were identified using a Grubbs’ test (alpha = 0.05) and normal distribution was assessed using the Shapiro-Wilk test. Variance was analyzed using a two-tailed Student’s t-test. Data is reported as means ± standard error of the mean (SEM), and significant differences were defined as a statistical threshold of *p* < 0.05.

## 3. Results

### 3.1. Atmospheric oxygen (21%) induces features of syncytiotrophoblast stress and injury relative to physiological (8%) oxygen conditions

To establish an appropriate physiological control condition, RCB 4940 cells were differentiated into STs and either maintained under atmospheric oxygen conditions (21% O_2_) or exposed to 8% O_2_. Compared with cells exposed to 8% O_2_, atmospheric oxygen induced several characteristic features of PE-like ST dysfunction. Levels of cell death were assessed in STs following 24 h of differing oxygen exposures through Annexin V/PI staining. Cells maintained at 21% O_2_ exhibited significantly increased cell death relative to cells exposed to 8% O_2_ (*p* < 0.0011), as determined by increased fluorescence of Annexin V and PI relative to total DAPI fluorescence, compared to cells exposed to 8% O_2_ indicating greater cell death under atmospheric oxygen conditions (**Fig. 1A-B**). To determine whether 8% O_2_ activated hypoxic signaling, HIF-1α (hypoxia-inducible factor 1-alpha) protein expression was assessed following 24 h of exposure. No significant difference in HIF-1α protein expression was observed between the two oxygen conditions (**Fig. 1C**), indicating that 8% O_2_ did not induce detectable HIF-1α stabilization relative to atmospheric oxygen. Secreted protein levels of sFlt-1 were increased in cells maintained at 21% O_2_ relative to cells exposed to 8% O_2_ following 24 h of exposure (Fig. 1D, *p* < 0.0001). Likewise, following 48 h of exposure, cells maintained at 21% O_2_ exhibited significantly increased *sFlt-1* (soluble fms-like tyrosine kinase-1) mRNA expression (Fig. 1E, *p* = 0.0007) and reduced *PlGF* (placental growth factor) mRNA expression (Fig. 1F, *p* = 0.0139) relative to 8% O_2_ exposure, indicative of a shift towards an anti-angiogenic phenotype. These findings demonstrate that atmospheric oxygen promotes an anti-angiogenic profile and increased cell death relative to 8% O_2_. As such, 8% O_2_ was selected as the physiological control condition for subsequent experiments.

**Fig. 1.**
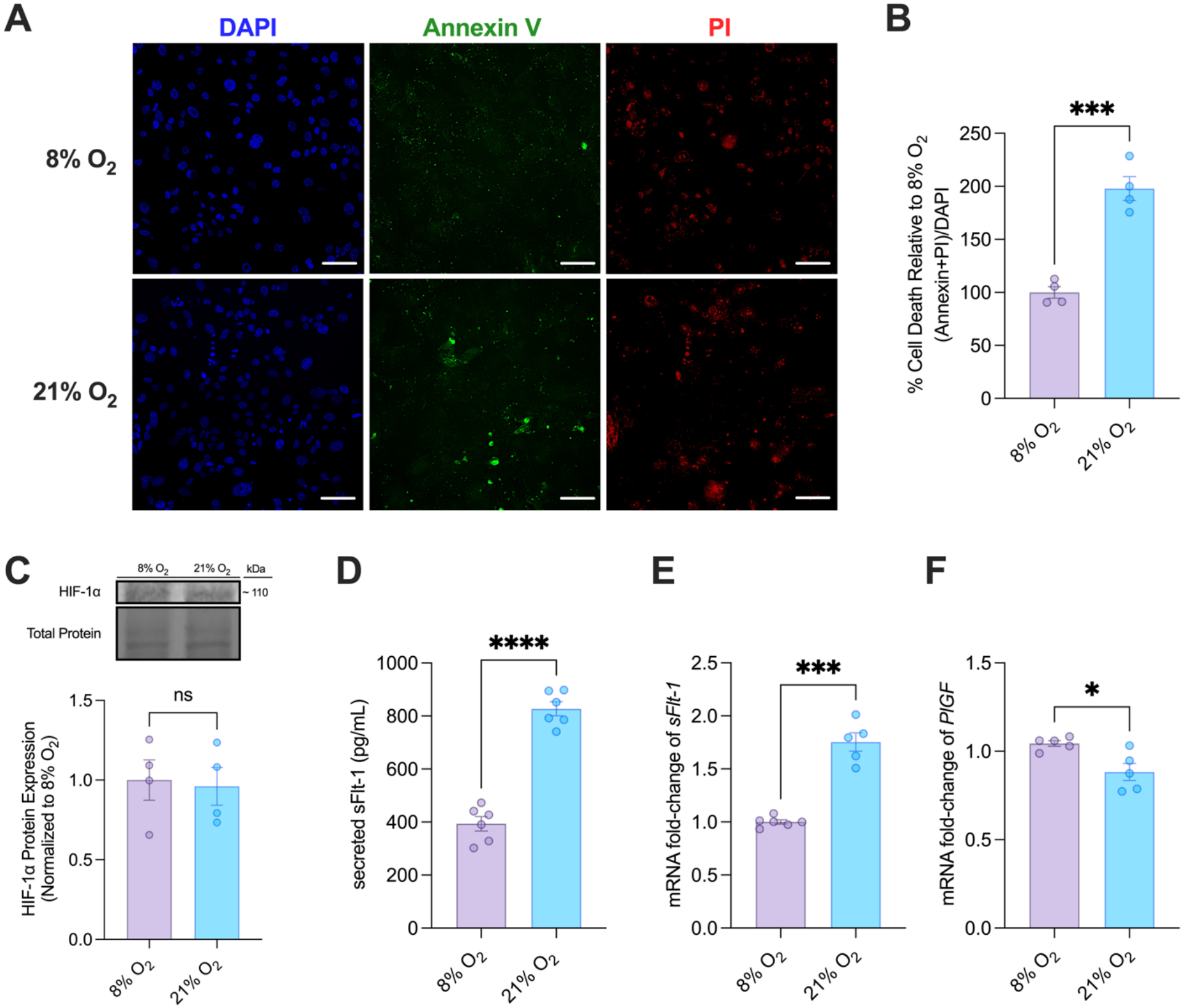
Atmospheric oxygen induces syncytiotrophoblast dysfunction relative to physiological oxygen conditions. Differentiated (ST) RCB 4940 cells were exposed to physiological oxygen (8% O_2_) or maintained at atmospheric oxygen (21% O_2_) for 24 h (A-D) or 48 h (E-F). **(A)** Representative fluorescence images corresponding to **(B)** quantification of cell death (Annexin and PI fluorescence/DAPI fluorescence). Cells were stained with Annexin V, Propidium Iodide (PI) and DAPI, fixed, and imaged using fluorescence microscopy on 1s exposure, 20X magnification (n=4). Scale bar = 100 µm. **(C)** Changes in protein levels of HIF-1α were normalized to stain-free images relative to 8% O_2_ controls, assessed using Western blotting (n=4). Cell lysates were separated by SDS-PAGE, and HIF-1α was detected using rabbit monoclonal anti-HIF-1α antibody. **(D)** Changes in secreted sFlt-1 protein levels were quantified in cell culture supernatants using ELISA (n=6). Protein concentrations (pg/mL) were calculated from standard curves generated for each assay. Changes in mRNA levels of *sFlt-1* **(E)** and *PlGF* **(F)** were normalized to *β-Actin* relative to 8% O_2_ controls, assessed using RT-qPCR (n=5-6). Significant differences were determined by a two-tailed Student’s t-test (\*\*\*\**p* < 0.0001, \*\*\**p* < 0.001, \*\**p* < 0.01, \**p* < 0.05). Data presented as mean ± SEM.

### 3.2. Acute and prolonged hypoxia induce distinct PE-associated angiogenic gene expression profiles in syncytiotrophoblasts

Having established 8% O_2_ as the physiological control condition, we next assessed how the duration of severe hypoxic exposure influenced PE-associated angiogenic marker expression. STs were exposed to hypoxic conditions (1% O_2_) for either 24 or 48 h and compared with cells maintained at 8% O_2_ for the corresponding duration. Following 24 h of exposure, STs exposed to 1% O_2_ exhibited a robust PE-like angiogenic gene expression profile. Compared to cells maintained at physiological oxygen conditions (8% O_2_), hypoxic cells exhibited increased mRNA of expression of *sFlt-1* (Fig. 2A, *p* < 0.0001), *sEng* (soluble endoglin; Fig. 2C, *p* < 0.0001), and *HIF1A* (Fig. 2D, *p* < 0.0001), along with reduced mRNA expression of *PlGF* (Fig. 2B, *p* < 0.0001), consistent with expression changes seen in PE. Notably, *sFlt-1* expression increased by over 60-fold following 24 h of 1% O_2_ exposure. In contrast, 48 h of 1% O_2_ exposure resulted in a distinct transcriptional response. While *sFlt-1* mRNA expression remained significantly increased in the hypoxic group (Fig. 2E, *p* < 0.0001), the magnitude of induction was lower than that observed in the 24 h exposure group. *PlGF* expression remained reduced after 48 h (Fig. 2F, *p* < 0.0001), and, unlike in the 24 h exposure group, mRNA expression of both *sEng* (Fig. 2G, *p* < 0.0001) and *HIF1A* (Fig. 2H, *p* = 0.0001) was significantly reduced after 48 h of hypoxic exposure. These findings demonstrate that the angiogenic transcriptional response to severe hypoxia varies with exposure duration.

**Fig. 2.**
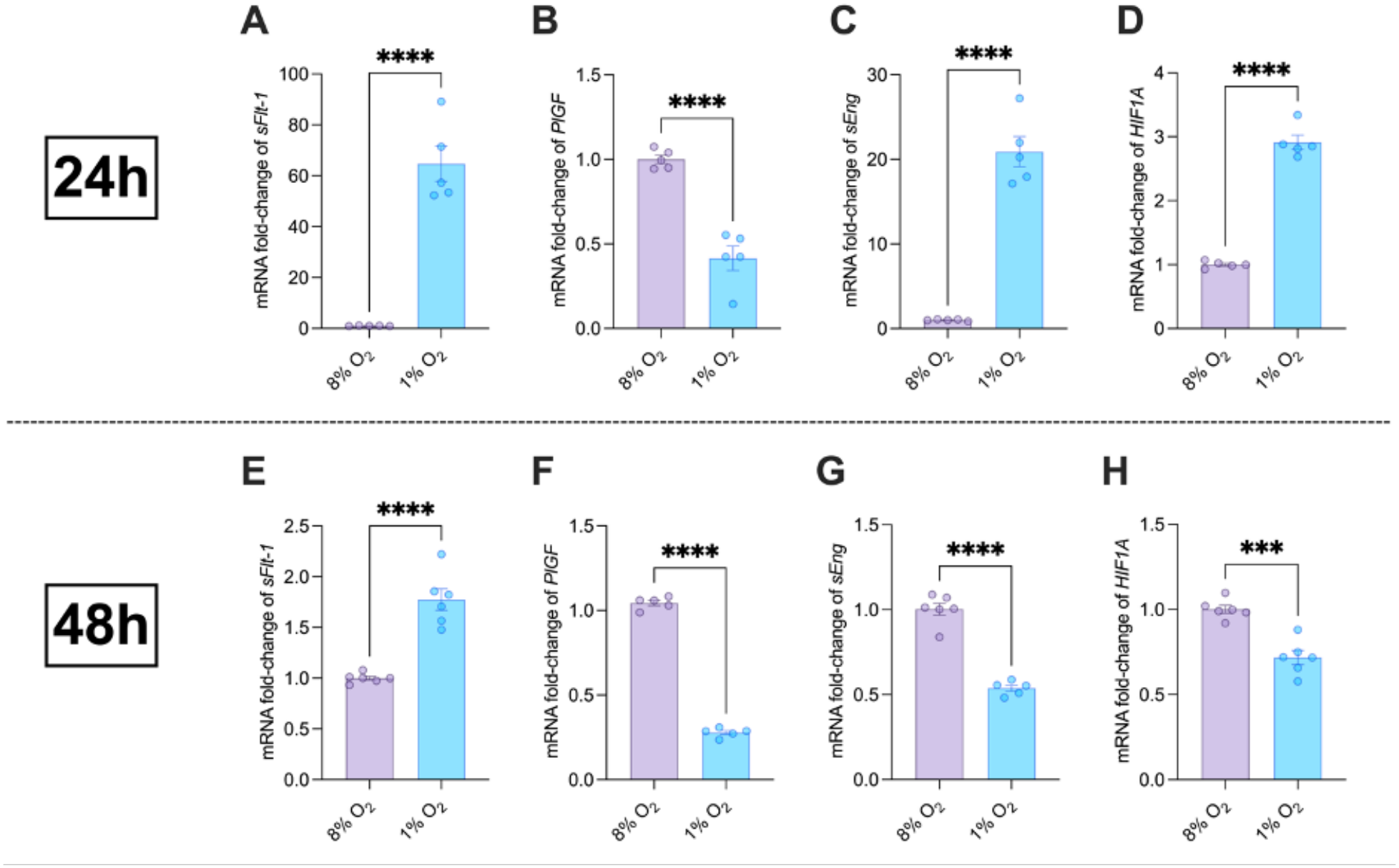
Prolonged hypoxia attenuates the acute induction of several PE-associated angiogenic genes. Differentiated (ST) RCB 4940 cells were exposed to 8% O_2_ or 1% O_2_ for 24 h (A-D) or 48 h (E-H). Changes in mRNA levels of *sFlt-1* **(A, E)**, *PlGF* **(B, F)**, *sEng* **(C, G)**, and *HIF1A* **(D, H)** were normalized to *β-Actin* relative to 8% O_2_ controls, assessed using qPCR. Significant differences were determined by a two-tailed Student’s t-test (\*\*\*\**p* < 0.0001, \*\*\**p* < 0.001). Data presented as mean ± SEM (n=5-6).

### 3.3. Hypoxia induces persistent changes in PE-associated angiogenic marker protein expression

To determine whether transcriptional changes in angiogenic markers were reflected at the protein level, secreted sFlt-1 and PlGF concentrations were quantified in cell culture supernatants by ELISA, while HIF-1α protein expression was assessed by Western blotting following 24- and 48-h hypoxic exposure. Consistent with the corresponding mRNA findings, 24 h of hypoxia (1% O_2_) significantly increased sFlt-1 concentrations (Fig. 3A, *p* < 0.0001), reduced PlGF concentrations (Fig. 3B, *p* < 0.0001), and increased HIF-1α protein expression (Fig. 3C, *p* = 0.0002) relative to physiological oxygen conditions. These changes persisted following prolonged hypoxic exposure. After 48 h, sFlt-1 concentrations remained significantly elevated (Fig. 3D, *p* < 0.0001), PlGF concentrations remained significantly reduced (Fig. 3E, *p* < 0.0001), and HIF-1α protein expression remained significantly increased relative to controls (Fig. 3F, *p* = 0.0005). Although the overall angiogenic imbalance was maintained at both the 24 h and 48 h time points, the magnitude of individual marker responses differed over time. The hypoxia-induced increase in sFlt-1 was more pronounced following 24 h than 48 h of exposure (**Fig. 3A,D**; 3.0-fold vs. 2.4-fold), whereas suppression of PlGF became more pronounced following prolonged hypoxia (**Fig. 3B,E**; 1.7-fold vs. 3.1-fold). In contrast, HIF-1α protein expression was increased to a similar extent at both time points (**Fig. 3C,F**; 3.7-fold). These findings demonstrate that hypoxia produces a sustained and temporally dynamic anti-angiogenic protein expression profile.

**Fig. 3.**
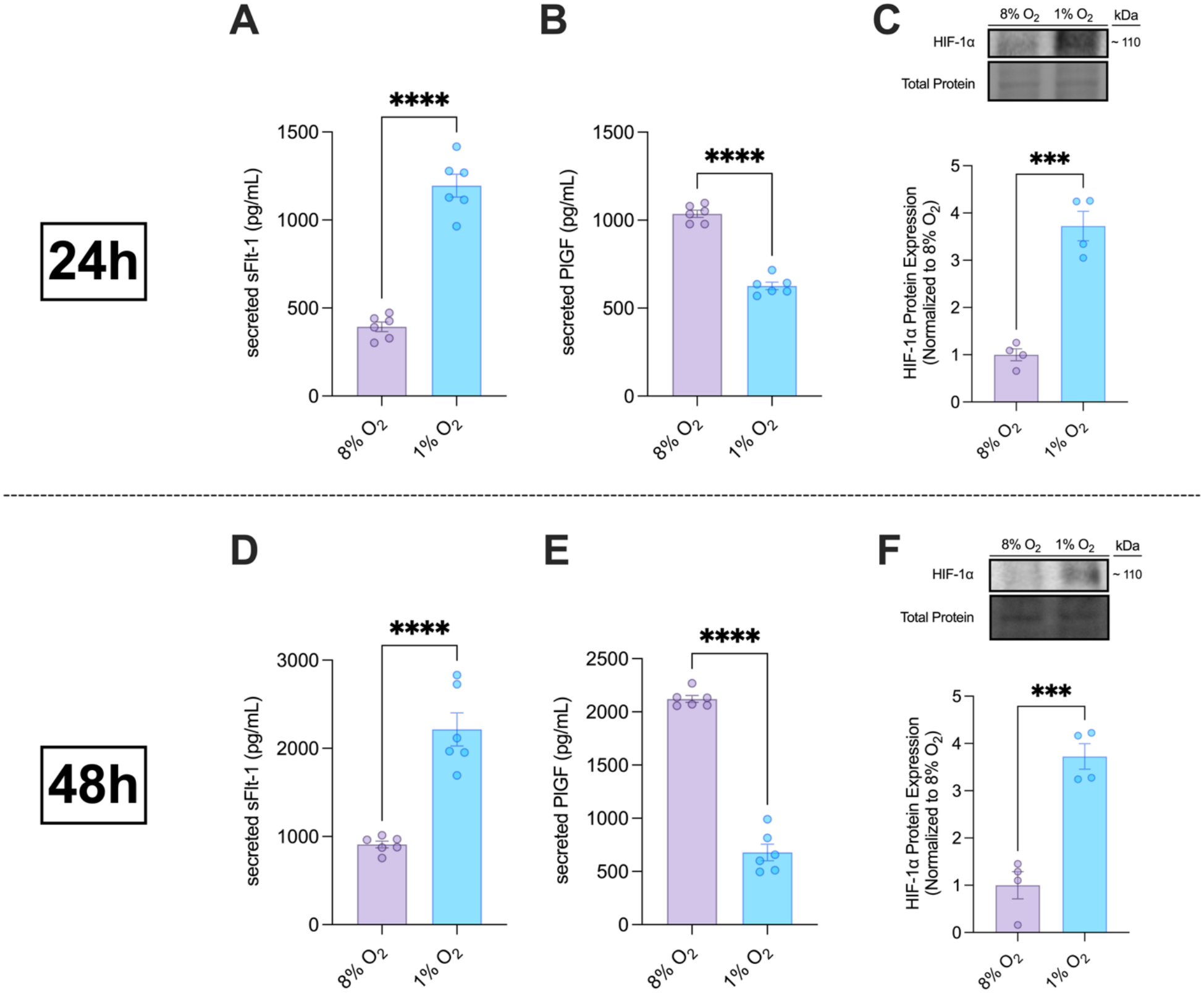
Both acute and prolonged hypoxia exposure reproduce PE-like angiogenic marker protein expression. Differentiated (ST) RCB 4940 cells were exposed to 8% O_2_ or 1% O_2_ for 24 h (A-C) or 48 h (D-F). Changes in secreted sFlt-1 **(A, D)** and PlGF **(B, E)** protein levels were quantified in cell culture supernatants using ELISA (n=6). Protein concentrations (pg/mL) were calculated from standard curves generated for each assay. **(C, F)** Changes in protein levels of HIF-1α were normalized to stain-free images relative to 8% O_2_ controls, assessed using Western blotting (n=4). Cell lysates were loaded and separated on SDS-PAGE, and content of HIF-1α was quantified using rabbit monoclonal anti-HIF-1α antibody. Significant differences were determined by a two-tailed Student’s t-test (\*\*\*\**p* < 0.0001, \*\*\**p* < 0.001). Data presented as mean ± SEM.

### 3.4. Acute, but not prolonged, hypoxia increases syncytiotrophoblast cell death

ST cell death following hypoxic exposure was assessed through Annexin V/PI staining. After 24 h, STs exposed to 1% O_2_ exhibited significantly greater apoptosis (Fig. 4B, *p* = 0.0023), necrosis (Fig. 4C, *p* = 0.0231), and overall cell death (Fig. 4D, *p* = 0.0002) than cells maintained under physiological oxygen conditions. In contrast, no significant differences in apoptosis, necrosis, or overall cell death were observed between oxygen conditions following 48 h of exposure (**Fig. 4F-H**). Apoptosis and necrosis were assessed by Annexin V and PI staining, respectively, relative to DAPI-positive nuclei, normalized to 8% O_2_. Overall cell death was quantified as the combined Annexin V and PI fluorescence normalized to DAPI-positive nuclei, normalized to 8% O_2_.

**Fig. 4.**
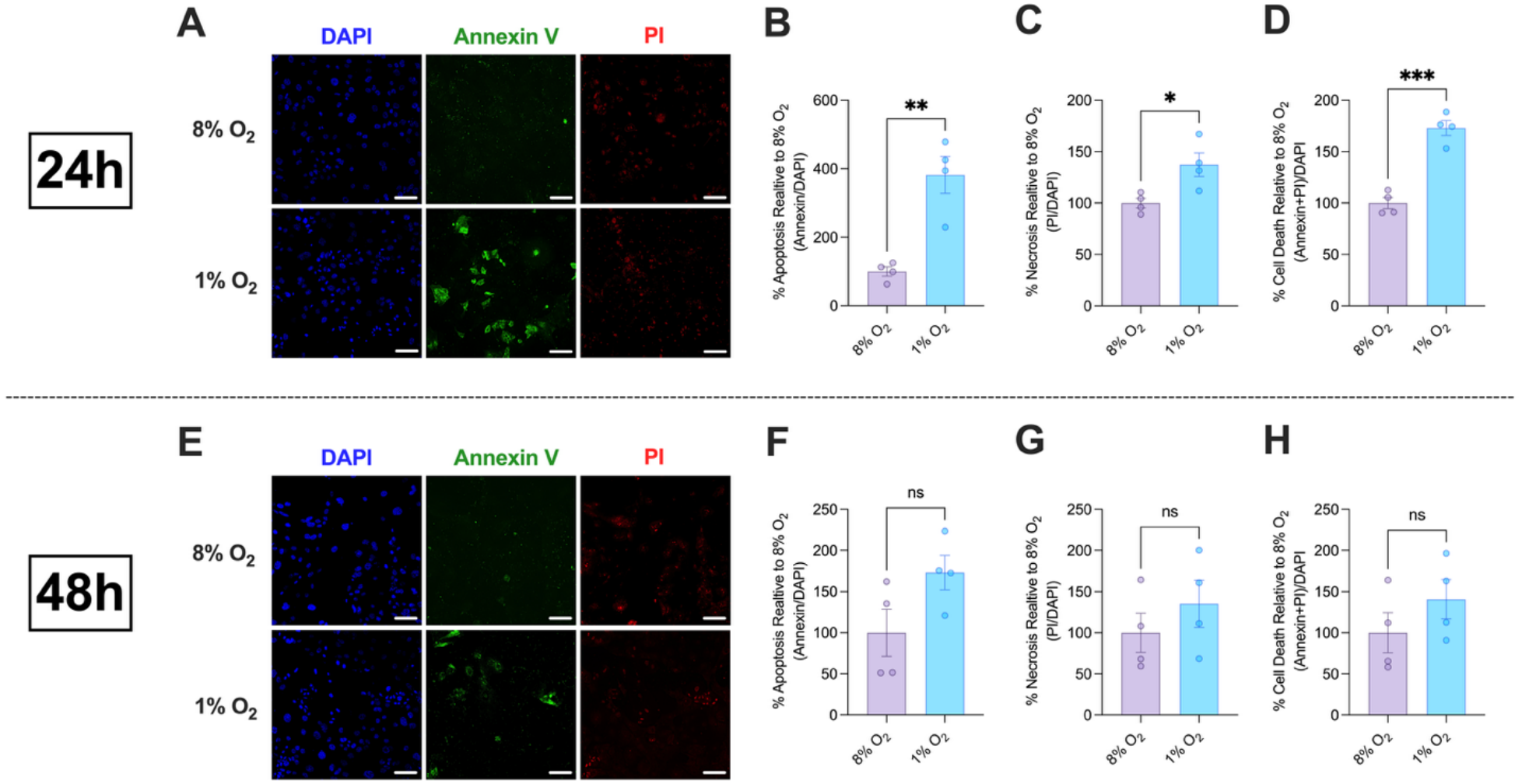
Acute hypoxia induces syncytiotrophoblast cell death, whereas prolonged hypoxia does not. Differentiated (ST) RCB 4940 cells were exposed to 8% O_2_ or 1% O_2_ for 24 h (A-D) or 48 h (E-H). **(A, E)** Representative fluorescence images corresponding to quantification of **(B, F)** apoptosis, **(C, G)** necrosis, and **(D, H)** overall cell death relative to 8% O_2_. Cells were stained with Annexin V, Propidium Iodide (PI) and DAPI, fixed, and imaged using fluorescence microscopy on 1s exposure, 20X magnification. Scale bar = 100 µm. Significant differences were determined by a two-tailed Student’s t-test (\*\*\**p* < 0.001, \*\**p* < 0.01, \**p* < 0.05). Data presented as mean ± SEM (n=4).

### 3.5. Acute and prolonged hypoxia differentially regulate mitochondrial stress and intrinsic apoptotic signaling in syncytiotrophoblasts

To further characterize the effects of hypoxia on ST cellular stress, markers of mitochondrial stress and intrinsic apoptosis were assessed following 24 and 48 h of exposure to 1% O_2_. After 24 h, hypoxia significantly increased mRNA expression of *BAX* (bcl-2-associated X protein; Fig. 5A, *p* = 0.0048), *VDAC1* (voltage-dependent anion channel 1; Fig. 5C, *p* < 0.0001) and *CASP9* (Caspase 9; Fig. 5C, *p* < 0.0001) relative to cells maintained under physiological oxygen conditions. Consistently, after 24 h of 1% O_2_, p53 protein expression was significantly increased (Fig. 5D, *p* = 0.0085), indicative of mitochondrial apoptotic stress pathway activation, and intracellular ROS levels were significantly increased (Fig. 5E, *p* = 0.0002), consistent with an acute oxidative stress response. In contrast, 48 h of 1% O_2_ exposure decreased mRNA expression of *BAX* (Fig. 5F, *p* < 0.0001), *VDAC1* (Fig. 5G, *p* = 0.0006) and *CASP9* (Fig. 5H, *p* = 0.0001) relative to physiological controls, while p53 protein and intracellular ROS levels were not significantly altered (**Fig. 5I-J**).

**Fig. 5.**
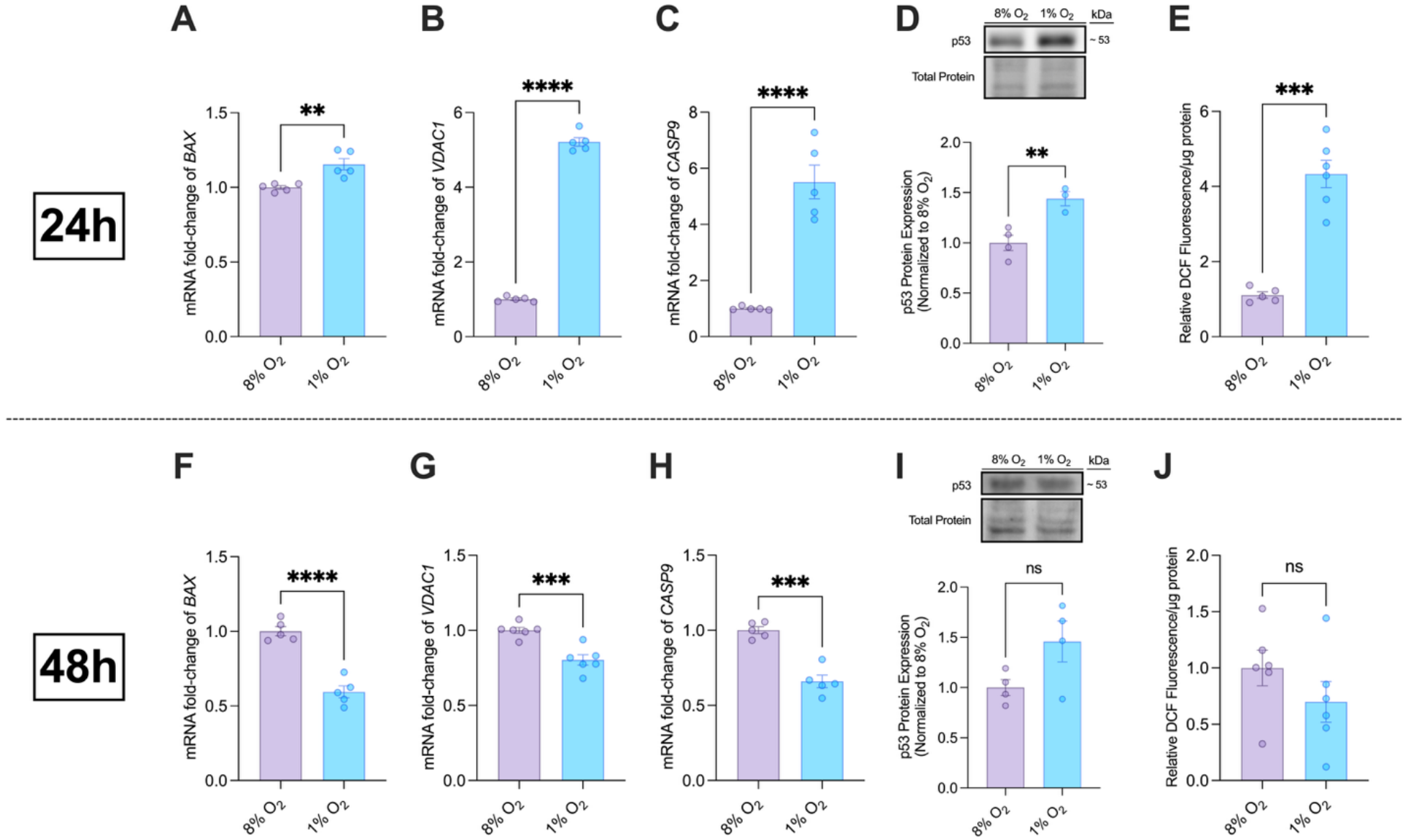
Acute hypoxia preferentially induces mitochondrial stress and intrinsic apoptotic signaling in syncytiotrophoblasts. Differentiated (ST) RCB 4940 cells were exposed to 8% O_2_ or 1% O_2_ for 24 h (A-E) or 48 h (F-J). Changes in mRNA levels of *BAX* **(A, F)**, *VDAC1* **(B, G)** and *CASP9* **(C, H)** were normalized to *β-Actin* relative to 8% O_2_ controls, assessed using RT-qPCR (n=5-6). **(D, I)** Changes in protein levels of p53 were normalized to stain-free images relative to 8% O_2_ controls, assessed using Western blotting (n=3-4). Cell lysates were separated on a SDS-PAGE, and content of p53 was quantified using rabbit polyclonal anti-p53 antibody. **(E, J)** The DCFDA assay was used to measure levels of reactive oxygen species relative to total protein in each well (n=5-6). Significant differences were determined by a two-tailed Student’s t-test (\*\*\*\**p* < 0.0001, \*\*\**p* < 0.001, \*\**p* < 0.01). Data presented as mean ± SEM.

## 4. Discussion

This study establishes an experimental framework for modelling hypoxia-induced syncytiotrophoblast dysfunction in vitro by defining a physiologically relevant oxygen control and characterizing the temporal effects of hypoxic exposure. Overall, our findings demonstrate that oxygen concentration and exposure duration critically influence trophoblast phenotype. Compared with 8% O_2_, atmospheric oxygen independently induced cellular stress and angiogenic dysregulation, indicating that the routine use of 21% O_2_ as a control may obscure or confound the interpretation of hypoxia-induced responses. In contrast, exposure to 1% O_2_ elicited a phenotype characterized by angiogenic imbalance, cell death, intrinsic apoptosis-associated signalling, and oxidative-stress related changes, with several responses being more pronounced following 24 h than 48 h of hypoxic exposure. These findings provide a framework for more physiologically accurate modelling of placental hypoxia and suggest that careful consideration of baseline oxygen conditions is essential for interpreting PE-associated trophoblast dysfunction in vitro.

A central finding of the present study was that atmospheric oxygen does not represent a biologically neutral control condition for differentiated syncytiotrophoblasts in the present model. Instead, culture at 21% O_2_ produced a subtle stress-associated phenotype relative to 8% O_2_, characterized by angiogenic dysregulation and increased cell death. This was characterized by increased transcript and secreted protein levels of sFlt-1, and decreased transcript levels of *PlGF* observed in the 21% O_2_ condition relative to 8% O_2_ (Figure 1). The angiogenic profile observed under 8% O2, reflected by sFlt-1 and PlGF expression, was more consistent with that reported in healthy, uncomplicated pregnancies, in which placental sFlt-1 levels are relatively lower and PlGF levels are higher than in PE [27]. In contrast, 21% O_2_ exposure resulted in increased sFlt-1 and reduced *PlGF* expression, characteristic of a shift toward an anti-angiogenic, PE-like phenotype [28,29].

The absence of a difference in HIF-1α protein expression between 8% and 21% O_2_ (Figure 1) suggests that 8% O_2_ does not induce a classical hypoxic response in our model and may instead reflect physiological oxygen conditions for STs [16,30]. Interestingly, these findings differ from our previous work in BeWo cells, in which differentiation under 8% O_2_ resulted in significantly increased HIF-1α protein expression relative to cells differentiated under atmospheric oxygen [21]. The discrepancy between these findings may reflect fundamental differences in trophoblast origin, differentiation state, and experimental design. Accordingly, physiological oxygen concentrations cannot be entirely generalized across trophoblast models and should instead be empirically established for the specific cell system under investigation.

Further supporting the physiological relevance of 8% O_2_ in our model, overall cell death was significantly greater under 21% O_2_ relative to 8% O_2_ (Figure 1). This is notable given that increased ST apoptosis and cell death are well-established features of PE pathology, suggesting that atmospheric oxygen conditions may induce a mild, non-physiological stress response in vitro [31–33]. Accordingly, 8% O_2_ was selected as the baseline condition for subsequent experiments.

Following the establishment of 8% O_2_ as a physiologically relevant baseline, we next examined how the duration of severe hypoxic exposure influenced the resulting ST phenotype. Considerable variability exists across the literature with respect to oxygen concentration, timing of oxygen exposure relative to trophoblast differentiation, method of hypoxia induction (e.g., hypoxic incubators versus chemical mimetics such as cobalt chloride), and exposure duration, with treatment periods ranging from several hours to multiple days [16,34]. Rather than producing a uniformly progressive response, 1% O_2_ elicited temporally distinct cellular states, whereby acute hypoxic exposure (24 h of 1% O_2_) generated a PE-associated stress phenotype, whereas several responses were either attenuated or sustained through distinct mechanisms following prolonged hypoxic exposure (48 h of 1% O_2_). These findings identify that exposure duration may be a critical determinant of the nature, rather than severity of hypoxia-induced ST dysfunction.

Overall, 24 h exposure to 1% O_2_ more closely reproduced the angiogenic phenotype associated with PE than prolonged exposure. We observed induction of *sFlt-1* and *sEng* expression following acute hypoxia, which is consistent with pronounced vascular and maternal endothelial dysfunction in PE, considered hallmark indicators and predictive biomarkers of the disease [35] (Figure 2). Although *sFlt-1* remained elevated after 48 h, its induction was attenuated, while *sEng* expression fell below physiological control levels. Likewise, secreted sFlt-1 concentrations increased to a greater extent following acute hypoxic exposure compared to prolonged exposure, although the attenuation observed following 48 h was considerably less pronounced than that observed at the mRNA level (Figure 3). This suggests that sustained hypoxia does not simply amplify the acute angiogenic response, but may instead engage adaptative or regulatory processes that limit the production of selected anti-angiogenic mediators. Of interest, our sFlt-1 findings align with those from a study that assessed secreted sFlt-1 expression following varying durations of oxygen deprivation in several trophoblast cell lines, where 12 and 24 h of hypoxia induced upregulation in secreted protein levels in BeWo cells, whereas 48 h of hypoxia did not [18].

sFlt-1 acts as a decoy receptor for pro-angiogenic factors, and as such limits blood vessel growth [36–38]. Similarly, sEng impairs angiogenesis by binding to transforming growth factor-β (TGF-β) family ligands, thus impairing vascular development [37–39]. Under physiological placental conditions, these anti-angiogenic markers help regulate the balance of placental vascular growth and development [40,41]. However, excessive production and release of sFlt-1 and sEng from the placenta into maternal circulation, as seen in PE, leads to endothelial dysfunction and contributes to the characteristic maternal vascular symptoms seen in the disease [42,43]. Interestingly, our study demonstrated that following 48 h of hypoxic exposure, STs exhibited reduced *sEng* expression relative to physiological controls, suggesting that prolonged hypoxia may drive adaptive responses that suppress the initial anti-angiogenic response.

*HIF1A* gene expression was dysregulated in a similar manner, with acute hypoxic exposure increasing mRNA expression and reduced gene expression following prolonged exposure (Figure 2). Given the central role of HIF-1α in coordinating cellular adaptation to hypoxia and promoting oxidative stress and anti-angiogenic signalling in PE, it is not surprising that increased HIF-1α expression has been consistently reported in PE placentas [44–46]. Notably, previous work has proposed that HIF-1α stabilization represents an acute response to severe hypoxia that diminishes with prolonged oxygen deprivation [45,47]. Consistent with this concept, *HIF1A* mRNA expression was reduced relative to controls following 48 h of hypoxia in the present study (Figure 2). Interestingly, however, HIF-1α protein expression remained elevated to a similar extent following both acute and prolonged hypoxic exposure (Figure 3). Although HIF transcriptional activity can be regulated independently of protein stability through oxygen-dependent modulation of interactions with transcriptional co-activators, the functional significance of the divergent HIF-1α mRNA and protein responses observed here remains unclear [47,48]. Future studies assessing downstream HIF target gene expression, nuclear localization, and DNA-binding activity will be important to determine the functional consequences of these differing responses.

In contrast to the anti-angiogenic factors, hypoxia-induced suppression of PlGF was evident following both acute and prolonged exposure, and the reduction was even more pronounced after 48 h at both the mRNA and soluble protein level (Figures 2 and 3). This response differed from that observed for sFlt-1, *sEng*, and *HIF1A*, whose induction was substantially attenuated following prolonged hypoxic exposure. This discrepancy in findings may suggest that pro- and anti-angiogenic factors exhibit distinct regulatory responses to sustained hypoxia. Consistent with this interpretation, previous studies have demonstrated that hypoxic regulation of PlGF and sFlt-1 occurs through distinct molecular mechanisms. Whereas hypoxia-induced sFlt-1 expression is mediated by HIF-1, PlGF regulation appears to occur independently of HIF-1 signalling [49,50].

Overall, the angiogenic profile observed following 24 h of hypoxia more closely resembles the characteristic imbalances seen in PE than that observed following prolonged hypoxic exposure, especially due to the robust induction of the key anti-angiogenic markers sFlt-1 and *sEng* [42,43]. In contrast, prolonged hypoxic exposure resulted in attenuation of *HIF1A* gene expression and reduced induction of anti-angiogenic factors, despite persistent HIF-1α protein expression, suggesting that additional regulatory mechanisms likely influence the angiogenic response during sustained hypoxia. Importantly, the continued suppression of PlGF following prolonged hypoxia further highlights that individual angiogenic mediators are regulated through distinct molecular mechanisms and exhibit different temporal responses to sustained oxygen deprivation [49,50]. As such, the suitability of a hypoxia-based PE model should be considered in the context of the specific biological pathways and the experimental question under investigation.

We next sought to characterize hypoxia-induced ST injury by assessing apoptosis, necrosis, and cell death together with markers of intrinsic apoptosis and mitochondrial stress. In PE, excessive ST apoptosis and cell death are well-established pathological features that contribute to placental dysfunction and increased shedding of trophoblast-derived apoptotic debris into the maternal circulation, thereby promoting endothelial dysfunction and maternal systemic responses characteristic of the disease [51–53]. Mitochondrial dysfunction is also increasingly recognized as a key contributor to PE pathogenesis and has been implicated in the activation of intrinsic apoptotic signaling [54–56]. Consistent with the angiogenic marker expression findings, hypoxia-induced trophoblast injury and mitochondrial stress were considerably more pronounced following 24 h of exposure than after 48 h.

In our model, acute hypoxia significantly increased apoptosis, necrosis, and overall cell death, accompanied by increased expression of the pro-intrinsic apoptotic mediators *BAX*, *Caspase 9* (*CASP9*), and p53 (Figures 4 and 5). This injury phenotype was further accompanied by a marked mitochondrial stress response, evidenced by increased *VDAC1* expression and elevated intracellular ROS levels (Figures 4 and 5). Collectively, these findings suggest activation of coordinated mitochondrial stress and intrinsic apoptotic signaling pathways that resemble the trophoblast dysfunction observed in PE. In contrast, these responses were markedly attenuated or absent following 48 h of hypoxic exposure, with no significant difference in apoptosis, necrosis, and cell death, along with *BAX*, *CASP9* and *VDAC1* expression falling below control levels, and normalization of p53 protein and ROS levels (Figures 4 and 5). These findings suggest that differentiated STs may undergo a degree of adaptation to sustained hypoxic stress, limiting activation of mitochondrial stress and apoptotic pathways despite continued oxygen deprivation.

Increased oxidative stress and mitochondrial dysfunction are well-established features of PE placentas and are thought to contribute to impaired trophoblast function and disease progression [57,58]. Elevated ROS can promote mitochondrial damage, while increased VDAC1 expression has been associated with altered mitochondrial membrane permeability and activation of intrinsic apoptotic signalling [57,59–61]. Accordingly, the concurrent induction of *VDAC1*, ROS, and pro-apoptotic mediators following acute hypoxia supports the presence of a mitochondrial stress program that closely resembles the mitochondrial dysfunction reported in PE placentas.

Interestingly, although hypoxia itself is known to induce mitochondrial dysfunction, oxidative stress and apoptosis in trophoblasts, elevated sFlt-1 has been suggested to drive these same pathological responses [13,62,63]. While this was not directly assessed in the present study, the less robust induction of sFlt-1 following prolonged hypoxia raises the possibility that attenuation of the anti-angiogenic response contributes, at least in part, to the reduced trophoblast injury observed after 48 h of exposure (Figures 2-6).

Taken together, the present findings demonstrate that acute (24 h) hypoxic exposure more consistently reproduces the molecular and cellular features of PE than prolonged (48 h) exposure. Across angiogenic and apoptotic signalling, ST injury/death, and mitochondrial stress, prolonged hypoxia was associated with attenuation or reversal of early pathological responses, suggesting that differentiated STs may undergo partial adaptation and/or selective cellular responses to sustained oxygen deprivation.

Importantly, the attenuation of pathological responses that was observed following prolonged hypoxic exposure should not be interpreted as evidence that longer-term hypoxic exposure lacks biological relevance. Instead, this may be indicative of distinct cellular mechanisms that occur following only prolonged oxygen deprivation. Although the precise mechanisms underlying this attenuation remain unclear, they may be driven by adaptive responses to prolonged hypoxia, alterations in hypoxia-responsive signalling, or selective survival of cells with greater resistance to hypoxic stress. While the present study demonstrates that acute, 24 h hypoxic exposure more closely reproduces several of the key features seen in PE, prolonged hypoxic exposure may provide valuable insight into the mechanisms of ST adaptation and chronic responses to oxygen deprivation. Furthermore, the ability of blastocyst-derived human trophoblast stem cells to be maintained as differentiated cultures for extended periods provides an opportunity to investigate the effects of longer durations of hypoxic exposure [24].

A major strength of this study is the systematic evaluation of both oxygen concentration and exposure duration using a physiologically relevant human blastocyst-derived ST model. This experimental framework provides an appropriate foundation for future in vitro studies while enabling direct comparison of physiological and pathological oxygen environments across multiple exposure durations.

An additional strength is the use of blastocyst-derived human trophoblast stem cells differentiated into STs, which more closely resemble human placental biology than commonly used transformed trophoblast cell lines [23–25,64]. Furthermore, the present study assessed several key PE-associated outcomes, including angiogenic factor imbalance, cell death, apoptosis, and mitochondrial stress, thus providing a broad assessment of hypoxia-induced ST dysfunction, rather than relying on a single disease endpoint. Overall, these findings establish a practical and physiologically relevant paradigm for modelling PE-associated ST dysfunction in vitro, while also providing considerations for selecting appropriate control and hypoxic exposure conditions in future studies.

Several considerations should be taken into account when interpreting these findings. First, while hypoxia is considered to be a central contributor in driving placental dysfunction and downstream clinical manifestations in PE, the disorder is multi-factorial, involving various other pathological processes such as inflammation, immune dysregulation, and endothelial dysfunction [6,65]. As such, hypoxia alone cannot fully reproduce the complexity of the in vivo disease environment [66,67]. Rather, the present study isolates the contribution of hypoxia to ST dysfunction under controlled experimental conditions. Additionally, the present study evaluated a selected panel of angiogenic, apoptotic, and mitochondrial markers. Although these endpoints capture several key features of PE-associated ST dysfunction, various additional molecular pathways have been implicated in PE and were beyond the scope of the present study [68,69].

Furthermore, the present study employed a two-dimensional monoculture model of differentiated STs. This approach enables investigation of ST-specific responses to hypoxia, but fails to account for interactions with other placental cell types, such as endothelial cells, extravillous trophoblasts, and maternal immune cells, which have all been shown to play a role in the pathophysiology of PE [70,71]. Future studies utilizing co-culture or placenta-on-a-chip models may help elucidate the role of multicellular interactions in a hypoxic placental environment and contextualize the findings of the present findings. Additionally, both early- and late-onset subtypes, which are thought to have overlapping but distinct underlying mechanisms [72]. Although impaired placentation and hypoxia are thought to play a particularly prominent role in EOPE, the present model is not intended to distinguish between PE subtypes and instead represents a reductionist approach for investigating hypoxia-induced ST dysfunction [10].

These findings emphasize the importance of selecting experimental conditions, including the timing and duration of oxygen exposure, according to the specific aspect of PE biology being investigated, as different oxygen paradigms may model distinct features of ST dysfunction. Although this model cannot reproduce the complex maternal clinical syndrome of preeclampsia, it provides a powerful platform for uncovering the mechanisms of hypoxia-induced placental dysfunction and advancing the identification of potential therapeutic targets.

## Author Contributions

J.S.F., S.L.W., S.R. conceptualization; J.S.F., S.R. methodology; J.S.F. investigation; J.S.F., C.M., T.P. data curation, J.S.F. formal analysis, J.S.F., S.L.W., S.R. funding acquisition, J.S.F., C.M., S.R. writing – original draft preparation, J.S.F., C.M., T.P., S.L.W., S.R. writing – reviewing and editing. All authors have read and approved the final version of the manuscript.

## Data Availability Statement

The data supporting the findings of this study are included within the article and its supplementary material. Further inquiries can be directed to the corresponding author.

## Acknowledgements

We thank Dr. Boyang Zhang and the Zhang Laboratory, McMaster University, for providing the human blastocyst-derived placental stem cell line used in this study.

