## Supplemental Figures S1-S4 and Table S1 for "Defining a hypoxic exposure paradigm for modelling preeclampsia-associated syncytiotrophoblast dysfunction in human blastocyst-derived placental stem cells"

### SUPPLEMENTARY INFORMATION

#### Supplementary Tables and Figures

**Table S1.** Primer sequences of human genes analyzed by RT-PCR

| **Gene** | **Forward (5′→3′)** | **Reverse (5′→3′)** |
|---|---|---|
| *sFlt-1* | CACACAGTGGCCATCAGC | CGATGACGATGGTGACGTT |
| *PlGF* | TGTCCAAAGTAGGGATGCGG | AGGCAACCACTGTTCTCCAG |
| *sEng* | GATCTCGATGCCGTCCTCAT | GGAGCTTGAAGCCACGAATG |
| *HIF1A* | ATTTTGGCAGCAACGACACA | CGTTTCAGCGGTGGGTAATG |
| *BAX* | GGGTTGTCGCCCTTTTCTAC | CGGAGGAAGTCCAATGTCCA |
| *CASP9* | GCCTGGACTTTCCTGCAAC | CCACTGTTCAGCACTTGTCG |
| *VDAC1* | GGGCAGTCTGGAAACCAAGTA | TAAACGTCAGGCCGTACTCAG |
| *β-actin* | TTACAGGAAGTCCCTTGCCATC | GCAATGCTATCACCTCCCCTG |

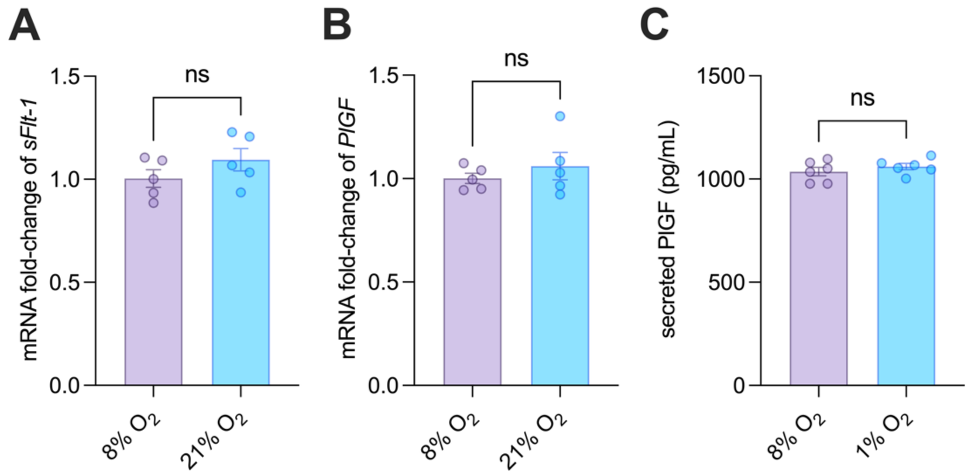


**Fig. S1. No significant differences were observed in select angiogenic markers following 24 h exposure to physiological and atmospheric oxygen conditions.** Differentiated (ST) RCB 4940 cells were exposed to physiological oxygen (8% $O_2$) or maintained at atmospheric oxygen (21% $O_2$) for 24h. Changes in *sFLT-1* **(A)** and *PlGF* **(B)** mRNA expression were normalized to *β-actin* relative to 8% $O_2$ controls and assessed by RT-qPCR (n=5-6). Secreted PlGF protein levels **(C)** were quantified in cell culture supernatants by ELISA (n=5-6). Significant differences were determined using a two-tailed Student's *t*-test. Data are presented as mean ± SEM.

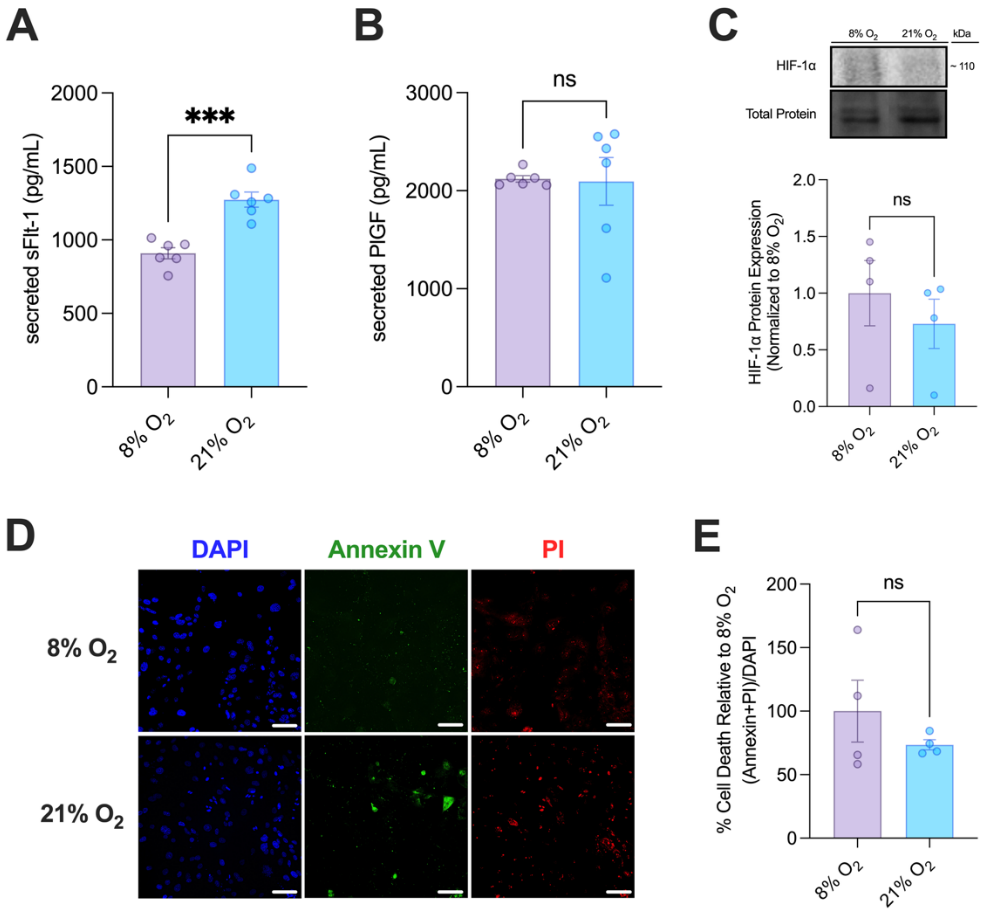


**Fig. S2. Select syncytiotrophoblast stress endpoints exhibit minimal changes following 48 h exposure to physiological and atmospheric oxygen conditions.** Differentiated (ST) RCB 4940 cells were exposed to physiological oxygen (8% $O_2$) or maintained at atmospheric oxygen (21% $O_2$) for 48h. Secreted sFlt-1 **(A)** and PlGF **(B)** protein levels were quantified in cell culture supernatants by ELISA (n=6). **(C)** HIF-1α protein expression was assessed by Western blotting and normalized to stain-free total protein relative to 8% $O_2$ (n=4). **(D)** Representative fluorescence images corresponding to **(E)** quantification of cell death (Annexin V + PI fluorescence normalized to DAPI) relative to 8% $O_2$ controls (n=4). Significant differences were determined by a two-tailed Student's t-test (***$p < 0.001$). Data presented as mean ± SEM.

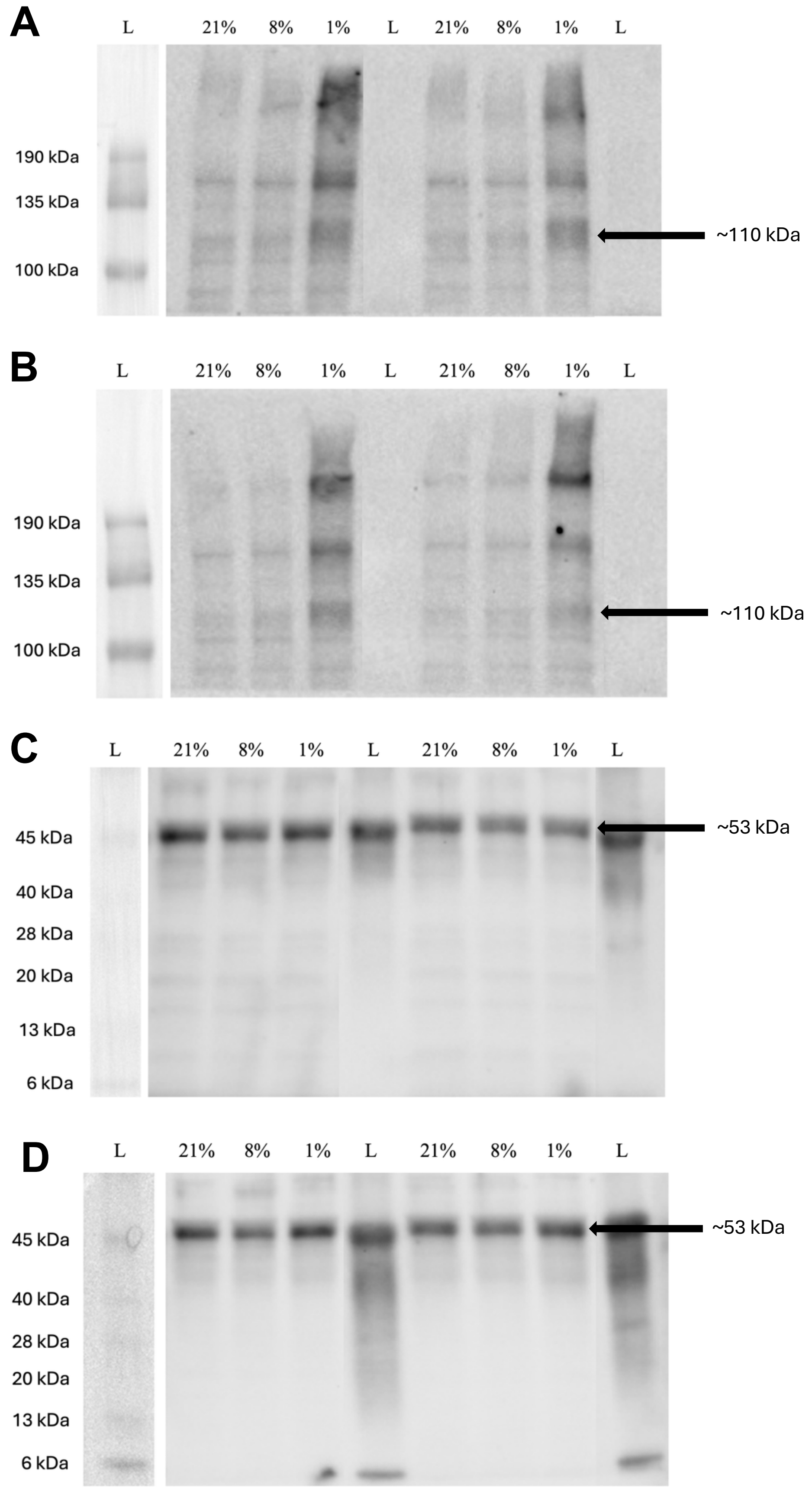
A
L 21% 8% 1% L 21% 8% 1% L
190 kDa
135 kDa
100 kDa
~110 kDa
B
L 21% 8% 1% L 21% 8% 1% L
190 kDa
135 kDa
100 kDa
~110 kDa
C
L 21% 8% 1% L 21% 8% 1% L
45 kDa
40 kDa
28 kDa
20 kDa
13 kDa
6 kDa
~53 kDa
D
L 21% 8% 1% L 21% 8% 1% L
45 kDa
40 kDa
28 kDa
20 kDa
13 kDa
6 kDa
~53 kDa

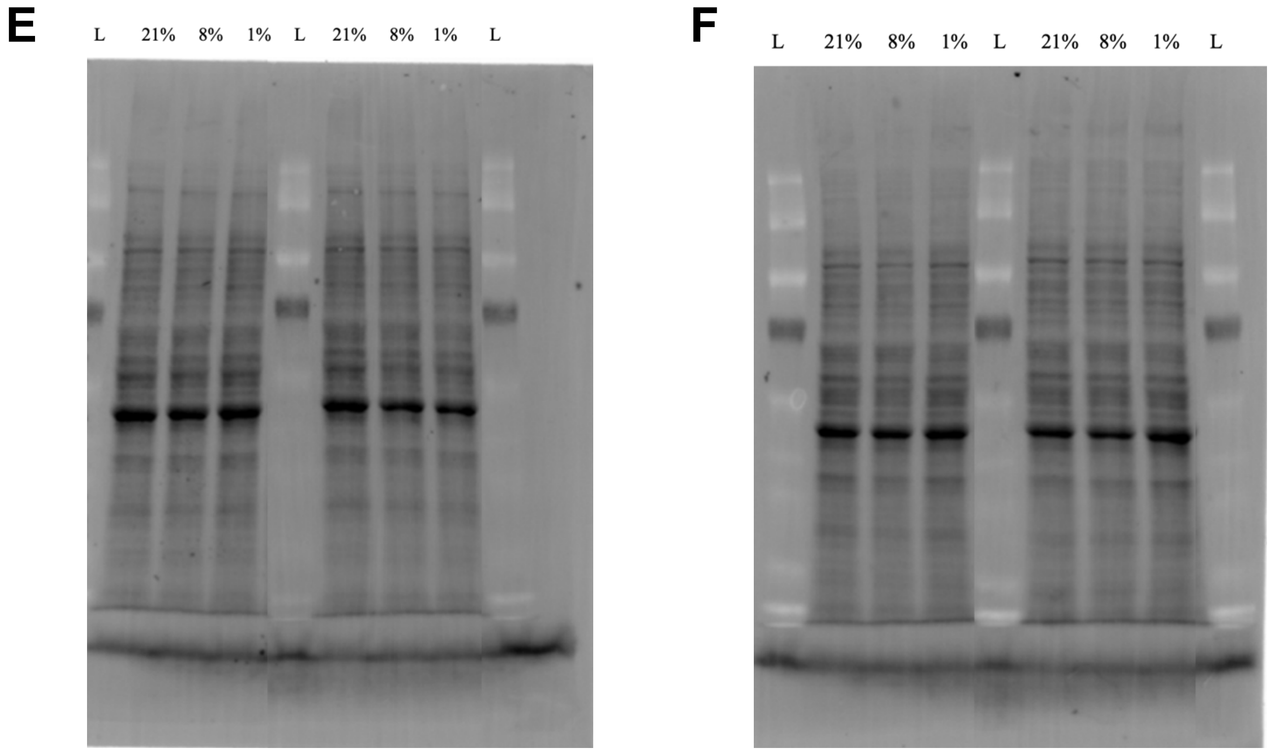


**Fig. S3. Full Western blot and corresponding stain-free gel images for the 24-hour exposure experiments.** Full Western blot images corresponding to **(A)** HIF1α (n1-n2); **(B)** HIF1α (n3-n4); **(C)** p53 (n1-n2); **(D)** p53 (n3-n4). Total protein visualized using Stain-Free gels and used for normalization for **(E)** HIF1α (n1-n2) & p53 (n1-n2); and **(F)** HIF1α (n3-n4) & p53 (n3-n4). Oxygen % of exposure group for the corresponding sample shown above each lane; molecular weight for protein band of interest denoted with an arrow; L = ladder. Blots were cut at ~70 kDa. Lanes corresponding to experimental conditions not included in the present study were omitted from the displayed membrane images.

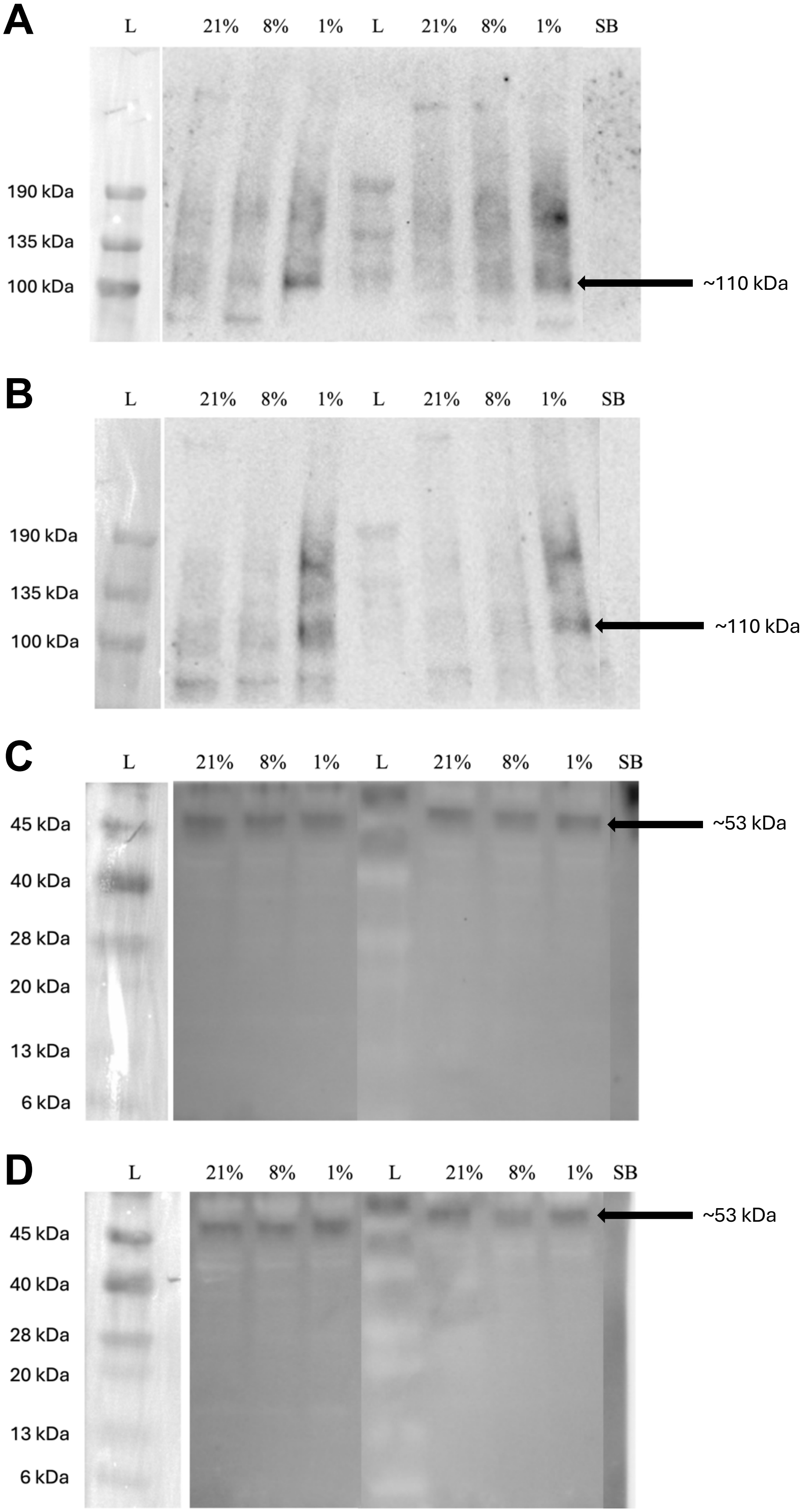
A
L 21% 8% 1% L 21% 8% 1% SB
190 kDa
135 kDa
100 kDa
~110 kDa
B
L 21% 8% 1% L 21% 8% 1% SB
190 kDa
135 kDa
100 kDa
~110 kDa
C
L 21% 8% 1% L 21% 8% 1% SB
45 kDa
40 kDa
28 kDa
20 kDa
13 kDa
6 kDa
~53 kDa
D
L 21% 8% 1% L 21% 8% 1% SB
45 kDa
40 kDa
28 kDa
20 kDa
13 kDa
6 kDa
~53 kDa

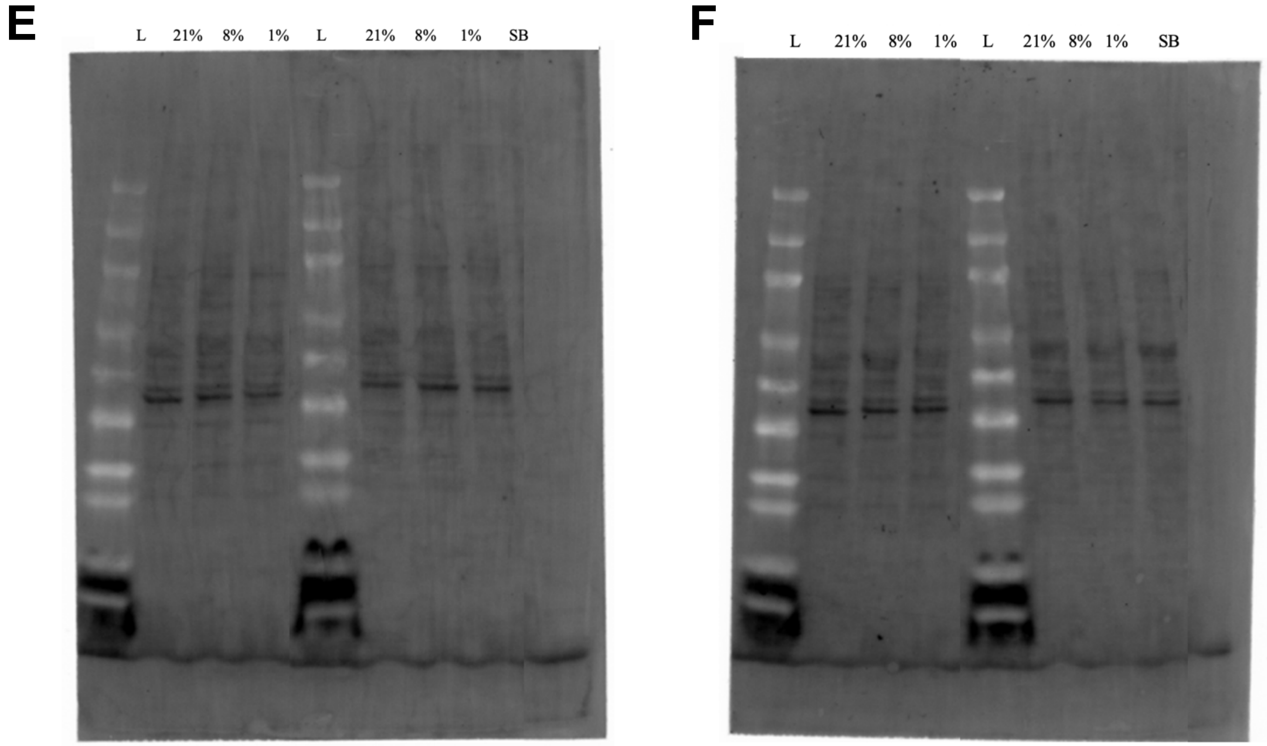


**Fig. S4. Full Western blot and corresponding stain-free gel images for the 48-hour exposure experiments.** Full Western blot images corresponding to **(A)** HIF1α (n1-n2); **(B)** HIF1α (n3-n4); **(C)** p53 (n1-n2); **(D)** p53 (n3-n4). Total protein visualized using Stain-Free gels and used for normalization for **(E)** HIF1α (n1-n2) & p53 (n1-n2); and **(F)** HIF1α (n3-n4) & p53 (n3-n4). Oxygen % of exposure group for the corresponding sample shown above each lane; molecular weight for protein band of interest denoted with an arrow; L = ladder, SB = sample buffer. Blots were cut at ~70 kDa. Lanes corresponding to experimental conditions not included in the present study were omitted from the displayed membrane images.
